# Enzyme-Regulated Non-Thermal Fluctuations Enhance Ligand Diffusion and Receptor-Mediated Endocytosis

**DOI:** 10.1101/2024.12.05.627033

**Authors:** Nividha, Arnab Maiti, Kshitiz Parihar, Rik Chakraborty, Pratibha Agarwala, Dibyendu K. Sasmal, Ravi Radhakrishnan, Dhiraj Bhatia, Krishna Kanti Dey

## Abstract

Active enzymes during catalyzing chemical reactions, have been found to generate significant mechanical fluctuations, which can influence the dynamics of their surroundings. These phenomena open new avenues for controlling mass transport in complex and dynamically inhomogeneous environments through localized chemical reactions. To explore this potential, we studied the uptake of transferrin molecules in retinal pigment epithelium (RPE) cells via clathrin-mediated endocytosis. In the presence of enzyme catalysis in the extracellular matrix, we observed a significant enhancement in the transport of fluorophore-tagged transferrin inside the cells. Fluorescence correlation spectroscopy measurements showed substantial increase in transferrin diffusion in the presence of active fluctuations. This study sheds light on the possibility that enzyme-substrate reactions within the extracellular matrix may induce long-range mechanical influences, facilitating targeted material delivery within intracellular milieu more efficiently than passive diffusion. These insights are expected to contribute to the development of better therapeutic strategies by overcoming limitations imposed by slow molecular diffusion under complex environments.

## Introduction

Enzyme catalysis plays a pivotal role in cellular function, facilitating critical processes like signalling and intercellular communication. (*1*) During catalytic reactions, enzymes not only diffuse to and from their substrates but also generate active fluctuations, a novel mechanobiological event which reflects the ability of enzymes to generate mechanical perturbations from chemical free energy. (*2*) In recent years, understanding the role of these fluctuations has become a key area of research in active matter physics. (*3–8*) Both theoretical (*9–14*) and experimental (*15–18*) studies have shown that the active fluctuations generated by enzymes can significantly influence the dynamics of their surroundings, even under artificially crowded conditions (*19*) that are usually characterized through diffusion enhancement of nearby passive tracers. This mechanistic behaviour, akin to that of ATP-driven molecular motors, suggests that enzymes may also function as force generators within cells, with potentially wide-ranging implications towards intracellular mechanics, opening an intriguing area for further research. These findings are particularly significant because molecular motors, known to drive diffusive, non-thermal motion of cellular components, play a major role in influencing the cell’s overall metabolic state. (*20, 21*) Similarly, the dynamic interactions observed between catalytic enzymes and their environment suggest that enzymes, beyond their primary role as catalysts, may act as mediators in complex dynamic interaction pathways. (*22*) Given that motor proteins operate in highly dissipative environments crowded with macromolecules, metabolites, and proteins (*23–25*), it is essential to investigate the implications of enzyme-generated fluctuations in similarly dense and crowded conditions. If enzymes can drive active fluctuations in such environments, this could have important implications for intracellular particle transport, which largely relies on passive diffusion, and motor protein facilitated motion. Enzyme-driven enhanced propulsion could also lead to significant advancements in the development of effective therapeutic strategies in complex biological environments. We investigated the effect of active fluctuations generated by enzymes in extracellular matrix over the transport of molecular ligands and their uptake in cells through clathrin-mediated endocytosis. Using Alexa Fluor 546 dye-tagged transferrin molecules (Tf-A546), as used in other studies, (*26*) we observed a striking 17% increase in transferrin uptake in retinal pigment epithelial (RPE1) cells in the presence of enzymatic activity in the extracellular environment. This activity-dependent ligand uptake was further substantiated by Fluorescence Correlation Spectroscopy (FCS) measurements, which revealed that the diffusion of Tf-A546 was enhanced by nearly 40% in the presence of enzyme catalysis in aqueous solutions. Although varying the ligand density in the extracellular environment increased the extent of endocytic uptake, internalization was significantly higher in the presence of active fluctuations across all concentrations of Tf-A546 used. Based on these observations, we hypothesized that enzyme-generated active fluctuations in the extracellular environment enhance transferrin diffusion, leading to their increased endocytic uptake in RPE cells. Enhanced diffusion of Tf-A546 is likely to facilitate more frequent ligand-receptor interactions at the cell membrane, accelerating pit formation and internalization. To confirm that this increase in transferrin uptake was indeed through clathrin-mediated endocytosis, we inhibited dynamin, a molecular scissor that regulates the internalization of clathrin-coated vesicles. In these experiments, no significant changes were observed in Tf-A546 uptake with or without enzyme activity, suggesting that enzyme-generated mechanical fluctuations primarily influenced the clathrin-mediated endocytic transport pathway. Our results suggest that the enzyme generated fluctuations offers opportunities to remotely control molecular diffusion and transport efficiencies in cells, especially for pathways like clathrin mediated endocytosis, where ligand internalization is largely governed by their diffusion towards the cell membrane. This study offers a fresh perspective on reaction controlled intracellular transport and its potential applications in targeted delivery (*27*), the mechanical behaviour of the extracellular milieu (*28*), and intracellular transport of molecules and metabolites. (*29, 30*) Our findings also provide new insights into how active fluctuations could significantly impact biochemical processes at the single particle level, offering exciting research opportunities in activity controlled cellular dynamics. (*31*)

## Results

We studied the effect of enzymatic activity on the cellular uptake of fluorescently labeled transferrin molecules (Tf-A546) shown in **Fig. 1**. For this, we used urease (from Canavalia ensiformis (Jack Bean), Sigma-Aldrich) and alkaline phosphatase (from bovine intestinal mucosa, Sigma-Aldrich) enzymes separately.These enzymes were selected because of their robustness and relatively high turnover rates at room temperature (urease: *k*_*cat*_ = 2.34 × 10^4^ s^−1^, alkaline phosphatase: *k*_*cat*_ = 1.4×10^4^ s^−1^. (*6,32*) They are also the enzymes whose enhanced diffusivity during catalysis has been established using different experimental techniques, such as fluorescence correlation spectroscopy (FCS) (*33*), dynamic light scattering (DLS) (*34*), optical microscopy followed by particle tracking (*35*) and total internal reflection fluorescence microscopy (TIRFM). (*36*) Transferrin (Tf-A546) molecules are used as markers for clathrin-mediated endocytosis (CME). The CME mechanism for transporting molecules across the cell membrane via a receptor protein (RP) is required to form clathrin-coated pits and vesicles (CCVs). (*37*) When the ligand molecules encounter the lipid membrane, RP binds with the ligand molecules (*38*), and subsequently, the RP-ligand complex interacts with clathrin through motifs at the clathrin terminal (*39*), forming CCV as shown in **Fig. 1**. In our studies, the cells were seeded 24 h before the experiment in Dulbecco’s Modified Eagle Medium (DMEM; Sigma-Aldrich) supplemented with 10% Fetal bovine serum (FBS; Gibco Make) and 1% penicillin/streptomycin (Pen-Strep; Gibco Make). For the experiment we incubate cells with Tf-A546 for 15 min at 37 °C in serum free DMEM media. The concentration of TfA546 used was 5 µg/mL and was kept constant in all the experiments. Without enzyme and the substrate, the treated cells were considered as control for validating the internalization efficiency of Tf-A546. For catalytic activity, the first enzyme we selected for our study was urease with its substrate urea. Urease converts urea into ammonium and bicarbonate ions. In experiments, we varied urease concentration between 0 – 5 nM, keeping substrate urea concentration constant at 100 mM. The internalization of Tf-A546 was characterized qualitatively and quantitatively using Confocal Laser Scanning Microscopy (CLSM), as shown in **Fig. 2**. The images were processed and quantified by using ImageJ (Fiji) software. Data normalization and statistical analysis were performed using GraphPad Prism 9.0. The data normalization process involved consideration of 0% as zero value and the average integrated density value of the control set as 100%, for all datasets. For statistical analysis, differences between the mean values of the control group with different test groups were calculated by using one-way ANOVA. Normalized fluorescence intensity per unit area (Φ) of Tf-A546 increased with increased enzyme concentration (keeping substrate amount constant), as shown in **Fig. 2c**. The enzyme and substrate concentrations were carefully selected to maintain constant reaction rates throughout the Tf-A546 uptake studies. We found that the normalized fluorescence intensity per unit area (Φ) of Tf-A546 in cells increased with increasing catalytic reaction and we calculated ∼ 17% enhancement corresponding to maximum catalytic activity. We confirmed that the increased fluorescence intensity was due to the catalytic activity by performing control experiments, as shown in **Fig. 2f**. For control experiments, we used only urea (100 mM) and only urease (5 nM) (without any reactions) keeping other conditions are same. We observed that the fluorescence intensity, measured in cells in the presence of enzyme activity was significantly different from the uptake of Tf-A546 in only media, indicating that the increased normalized fluorescence intensity was due to the enzymes’ activity during catalysis. To further confirm that the fluorescence intensity was not due to product formation during the urea-urease reaction (which increased the pH of the system), we performed a control experiment with Tf-A546 uptake in cells exposed to the product of the urea-urease reaction. The product was formed after a 15 min reaction with 5 nM urease and 100 mM urea. Again, we did not observe any significant changes in fluorescence intensity of Tf-A546 uptake, similar to the results obtained for water, only enzyme and only substrate solutions **Fig. 2f (A-D)**. These observations indicate that the enhanced uptake of transferrin was due to mechanical fluctuations generated by the enzymes during catalytic substrate turnover and was a function of the turnover rate.

**Figure 1:**
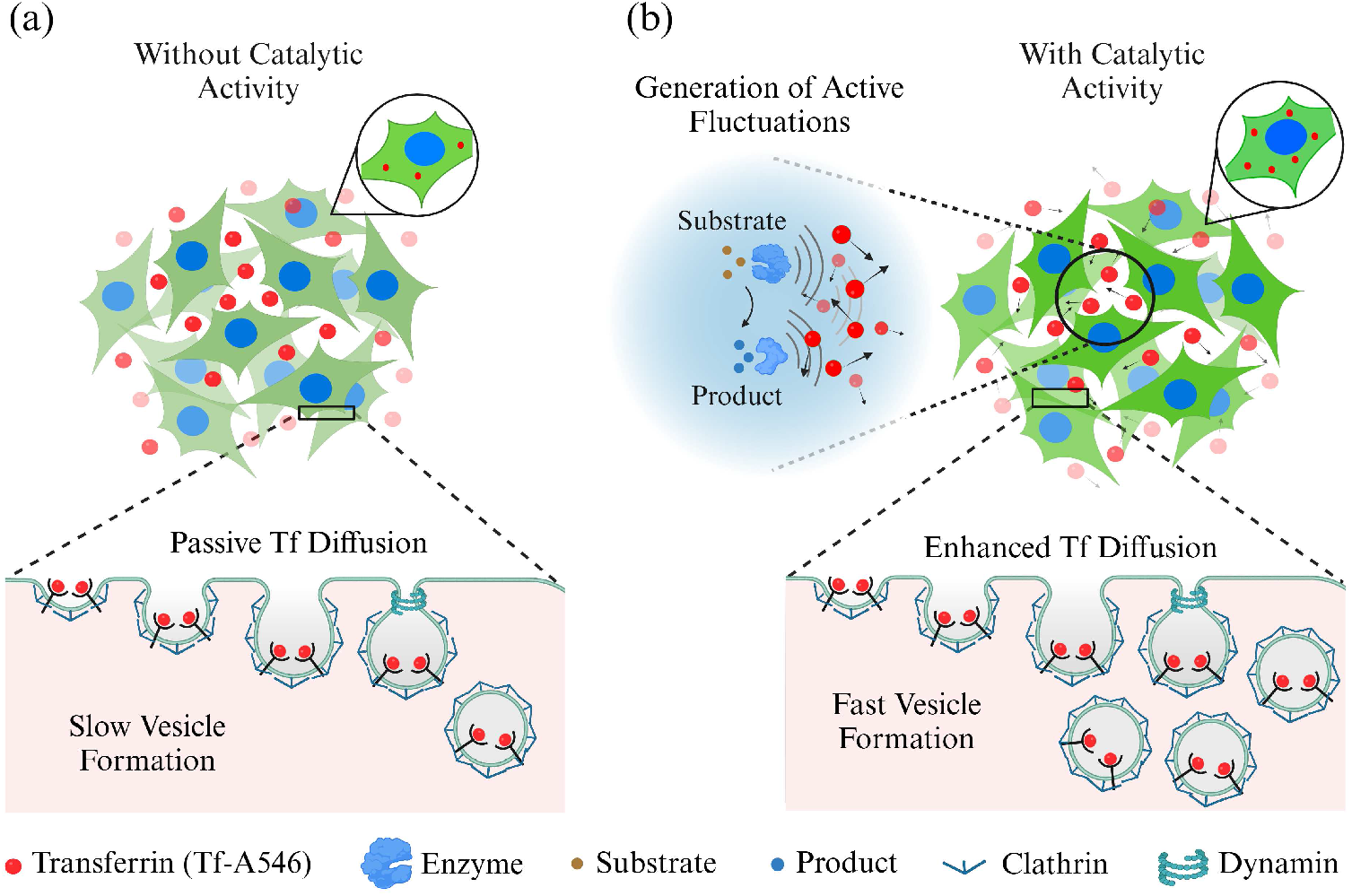
Effect of active fluctuations on Transferrin internalization. (**a**) In the absence of active fluctuations, passive diffusion of transferrin (Tf) molecules forms less ligand-receptor complex, leading to slower vesicle formation during endocytosis and less internalization of Tf. (**b**) In the presence of active fluctuations, diffusion of transferrin molecules is enhanced, leading to more ligand-receptor complex formation, which accelerates vesicle formation during endocytosis and increases internalization of Tf.

**Figure 2:**
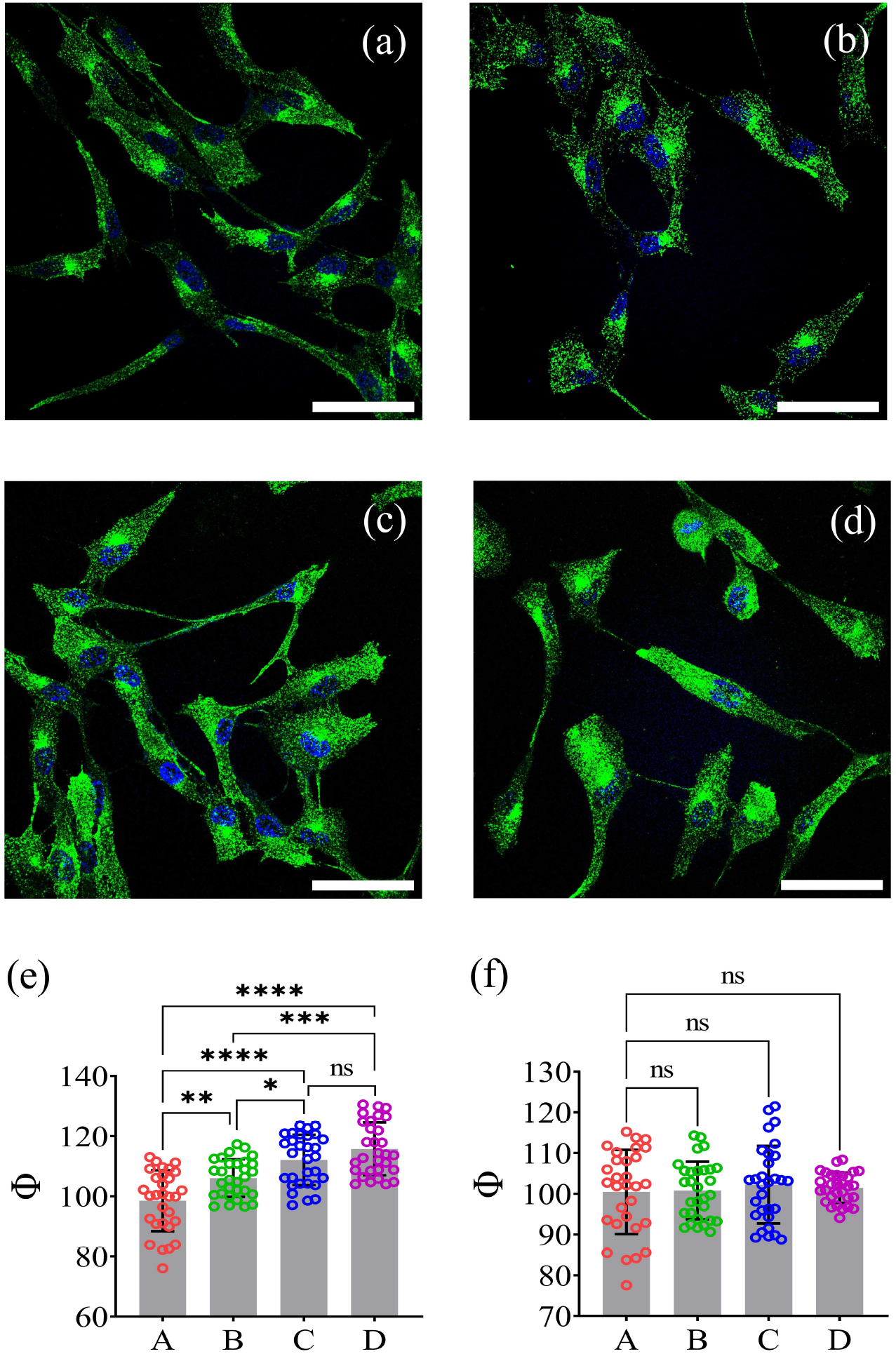
Uptake of Tf-A546 in RPE1 cells (shown in green) in the presence of urease activity. Tf-A546 uptake (a) without and (b-d) with enzymatic activity in the extracellular matrix. The urease concentrations in (b), (c), and (d) are 1 nM, 3 nM, and 5 nM, respectively, while the urea concentration was kept constant at 100 mM in each case. (f) Φ values for endocytic uptake of Tf-A546 in (A) media only, (B) 5 nM urease, (C) 100 mM urea, and (D) in the presence of urease-urea reaction products (prepared with 5 nM urease and 100 mM urea). The Φ values are calculated from data from 30 cells. Scale bars represent 60 µm and error bars represent the standard error of uptake across 30 cells. The symbols ∗, ∗∗, ∗∗∗ and ∗∗∗∗ indicate significance levels of p < 0.05, p < 0.01, p < 0.001 and p < 0.0001 respectively, while ns denotes not significant.

To test the generality of enzyme activity on cellular transport, we studied another enzyme, alkaline phosphatase (AKP), which catalyzes p-nitrophenyl phosphate (p-NPP) into p-nitrophenol (p-NP) and inorganic phosphate (P). The *k*_*cat*_ value of both urease and alkaline phosphatase is within the same order of magnitude. Therefore, it is expected that they will generate the same order of active fluctuations during substrate turnover. In experiments, we varied AKP concentration between 0 - 0.6 nM, keeping the substrate concentration constant at 1 mM. All other transferrin uptake protocol followed was the same as those in the case of experiments with urease. Here also we observed that the fluorescence intensity per unit area (Φ) of transferrin in cells increased with enzyme concentration. The maximum percentage increase in transferrin uptake for AKP was ∼ 17% (**Fig. 3e**) and here also we did not observe any significant change in fluorescence intensity of Tf-A546 in the presence of the individual reactants, as shown in **Fig. 3f**.

**Figure 3:**
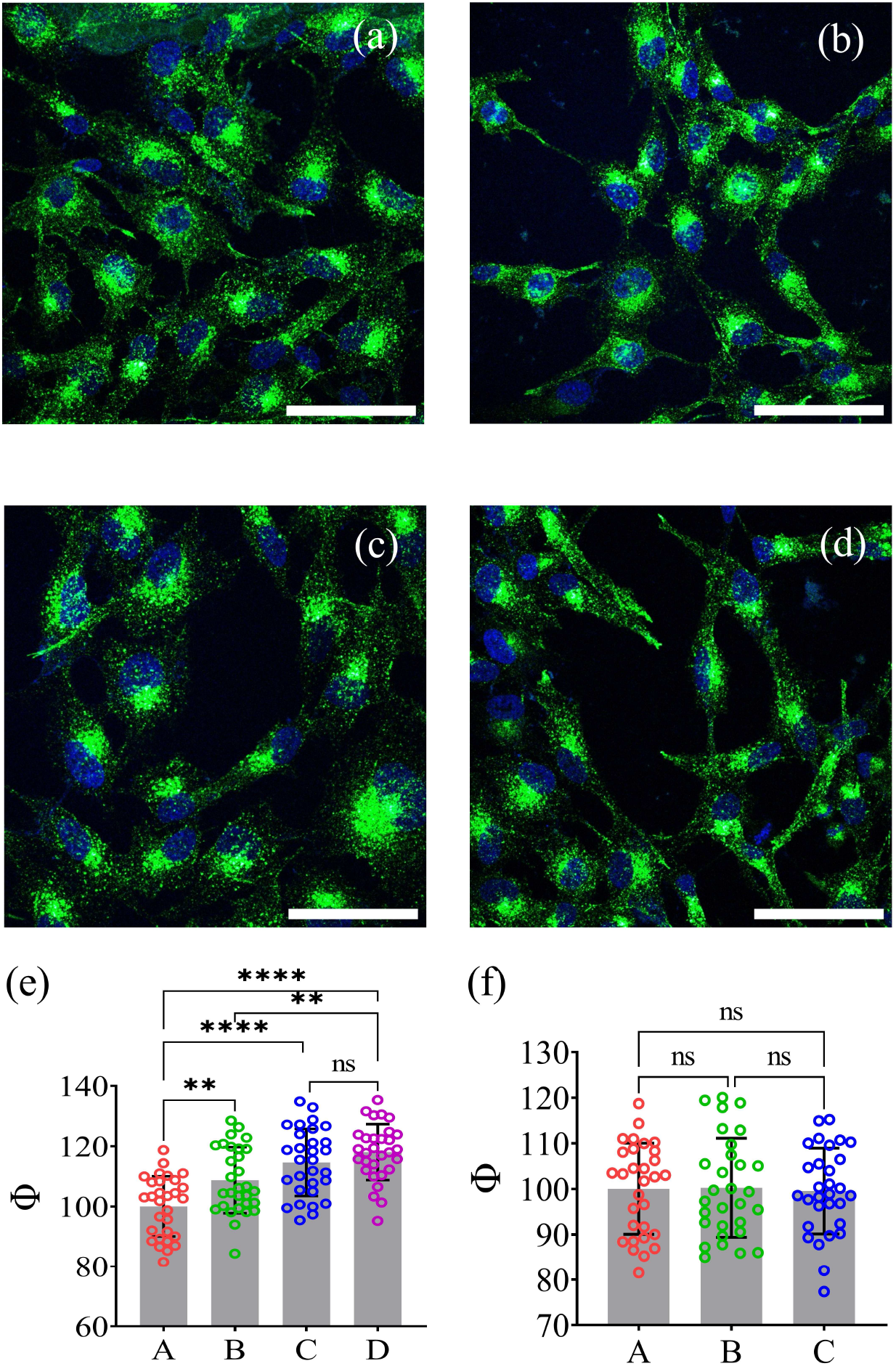
Uptake of Tf-A546 in RPE1 cells (shown in green) in the presence of alkaline phosphatase (AKP) activity. Tf-A546 uptake (a) without and (b-d) with enzymatic activity in the extracellular matrix. The AKP concentrations in (b), (c), and (d) are 0.2 nM, 0.4 nM, and 0.6 nM, respectively, while the p-NPP concentration was kept constant at 1 mM in each case. (e) Variation of Φ corresponding to Tf-A546 uptake in (A) media only, (B) 0.2 nM AKP and 1 mM p-NPP, (C) 0.4 nM AKP and 1 mM p-NPP, and (D) 0.6 nM AKP and 1 mM p-NPP. (f) Φ values recorded for endocytic uptake of Tf-A546 in (A) only media, (B) 0.6 nM AKP, (C) 1 mM p-NPP. The Φ values are calculated from data collected from 30 cells. Scale bars represent 60 µm and error bars represent the standard error of uptake across 30 cells. The symbols ∗∗ and ∗ ∗ ∗∗ indicate significance levels of p < 0.05 and p < 0.0001 respectively, while ns denotes not significant.

As studied by Mikhailov and Kapral, enzymes in cells may be assumed to behave as stochastic oscillating force dipoles while undergoing catalytic reactions, establishing hydrodynamic coupling with their surroundings. This coupling, in turn, could collectively impact the diffusion of nearby passive molecules. (*9, 10*) Validating the proposition, our previous studies have shown that active enzymes in solution could act as localized sources of mechanical energy that could significantly influence the diffusion of nearby passive molecules and microscopic particles both under dilute aqueous and artificial crowded environments. (*15, 18, 19*)

In the context of Tf internalization, changes in diffusion of the Tf can impact its binding with the Tf receptor (TfR) on the cell membrane. Specifically, the binding rate *k*_*a*_ strongly dependent on the collision frequency of Tf molecules to Tf receptors. The collision frequency, with TfR being stationary on the cell membrane, is predominately determined by the diffusion of Tf molecules in the solution. That is, enhanced diffusion of Tf can lead to more collisions with TfR which would then lead to higher probability of bond formation and subsequent internalization. To ascertain the effect of enzymatic activity on the diffusion of Tf-A546, we used fluorescence correlation spectroscopy (FCS). Details of FCS instrumentation, diffusion measurement procedure and data analysis protocols are provided in the Supplementary Materials (SM). In FCS studies, we observed that the diffusivity of Tf-A546 molecules increased with the increase in enzyme catalysis, indicating substrate turnover inducing active fluctuations and corresponding transfer of mechanical energy at the molecular scale. Here, we propose that the observed diffusion enhancement of the transferrin molecules was due to the generation of active fluctuations from active enzyme molecules to the surroundings. We measured diffusion of Tf-A546 in active enzyme solutions in the presence and absence of catalysis. We observed that the enhance diffusion of Tf-A546 is ∼ 40% in the presence of urease **Fig. 4c** and ∼44% in the presence of AKP activity respectively **Fig. 4d** We also performed control experiments to show that the individual reactants did not change the diffusion coefficient of the Tf-A546 without any catalysis (see **Figs. S3** and **S4** in the SM). Based on the FCS measurement results, we would thus expect that the receptor-ligand (Tf-TfR) binding rate at the cell membrane increased due to enhanced Tf diffusivity in the presence of active urease and AKP respectively. While the energetics of vesiculation for endocytosis has been extensively studied before using mesoscale models (*40*), here our primary focus was to delineate how changes in binding rate may influence internalization of Tf for which we used an ordinary differential equation (ODE) based model (see SM for details). Key parameters of the model are binding rate (*k*_*a*_), dissociation constant (*K*_*D*_), and internalization rate (*k*_*int*_). For different values of *K*_*D*_ and *k*_*int*_, we see that amount of Tf bound TfR internalized at 15 min (experimental time for our system) increases with increasing *k*_*a*_ up to *k*_*a*_ ∼ 0.005 *nM*^−1^ *min*^−1^ and plateaus for any further increase in *k*_*a*_ (see **Fig. 5 a-c**). This is consistent with what we observe in experiments where increasing concentration of active enzymes increases Tf internalization only up to certain concentration of enzyme (3 nM for urease (**Fig. 2e**) and 0.4 nM for AKP (**Fig. 3e**) after which Tf uptake plateaus. This suggests that higher enzyme activity may lead to higher diffusivities, as shown in **Fig. 4**, which would then lead to greater binding rate but after a certain value any further increase in binding rate would not lead to more internalization. This is because in our system the concentration of Tf-A456 was much higher than the number of TfR available (see SM), and TfR being the limiting reactant, constrains how much Tf-A456 can be internalized by cells. Overall, the experimental and model results suggests that the observed increase in Tf internalization in the presence of enzyme catalysis could, at least in part, be driven by enhanced diffusion leading to increased Tf-TfR binding rate.

**Figure 4:**
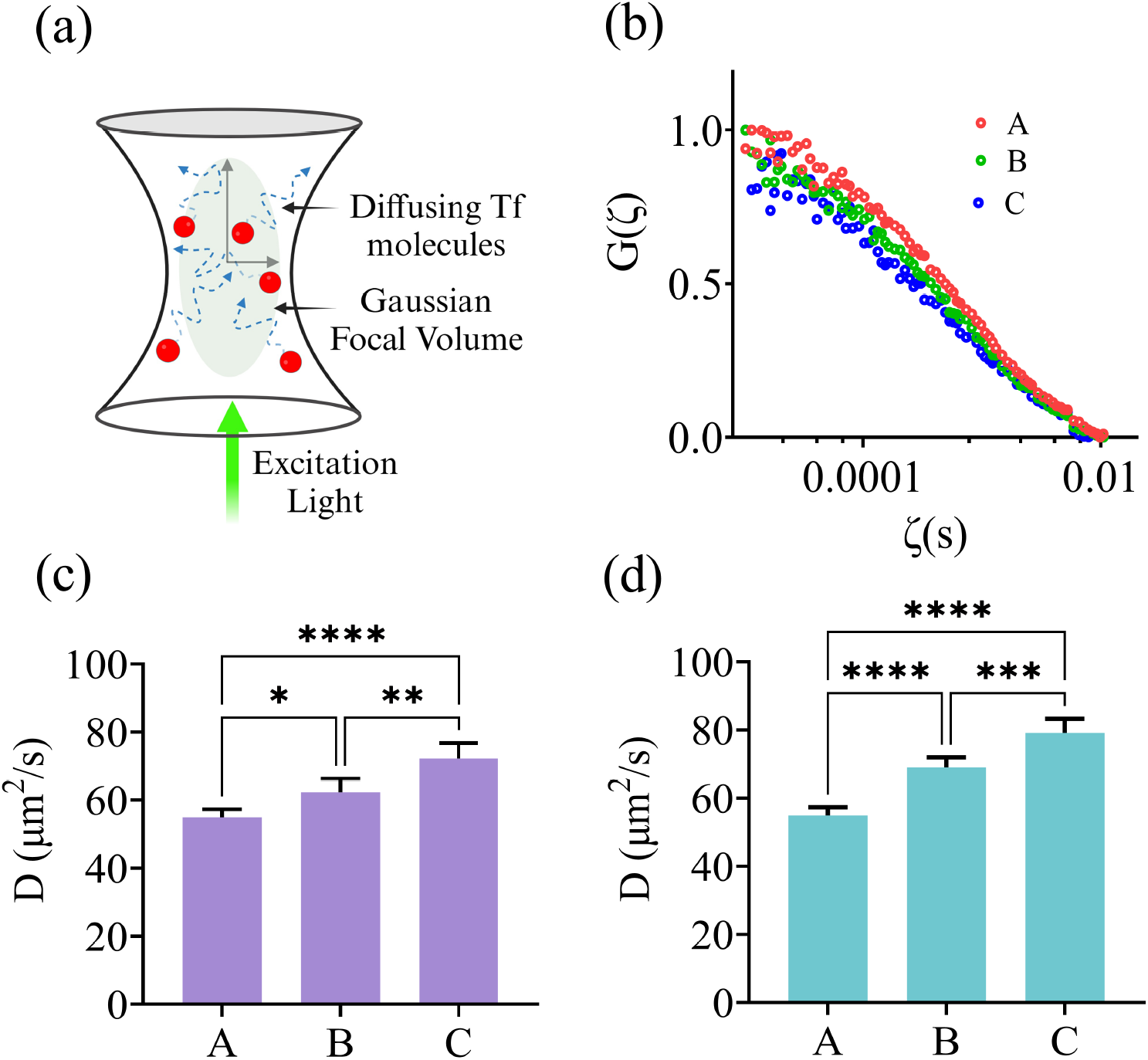
Diffusion measurement of Tf-A546 through Fluorescence Correlation Spectroscopy (FCS). (a) Schematic of diffusion of fluorescent molecules (Tf-A546) in and out of the confocal volume in FCS. (b) Autocorrelation function of diffusing molecules (Tf-A546) with diffusion time in (A) water, (B) 3 nM urease and 100 mM urea, (C) 5 nM urease and 100 mM urea. (c) Measured diffusion coefficients (D) of Tf-A546 in (A) DI water, (B) 3 nM urease and 100 mM urea, and (C) 5 nM urease and 100 mM urea. (d) Measured diffusion coefficients (D) of Tf-A546 in (A) DI water, (B) 0.4 nM AKP and 1 mM p-NPP, and (C) 0.6 nM AKP and 1 mM p-NPP. The diffusion coefficient of Tf-A546 was increased by up to 40% in the presence of active urease and by nearly 44% with active AKP. Error bars represent the standard deviations from three independent measurements under identical conditions. The symbols ∗, ∗∗, ∗∗∗ and ∗∗∗∗ indicate significance levels of p < 0.05, p < 0.01, p < 0.001 and p < 0.0001 respectively, while ns denotes not significant.

**Figure 5:**
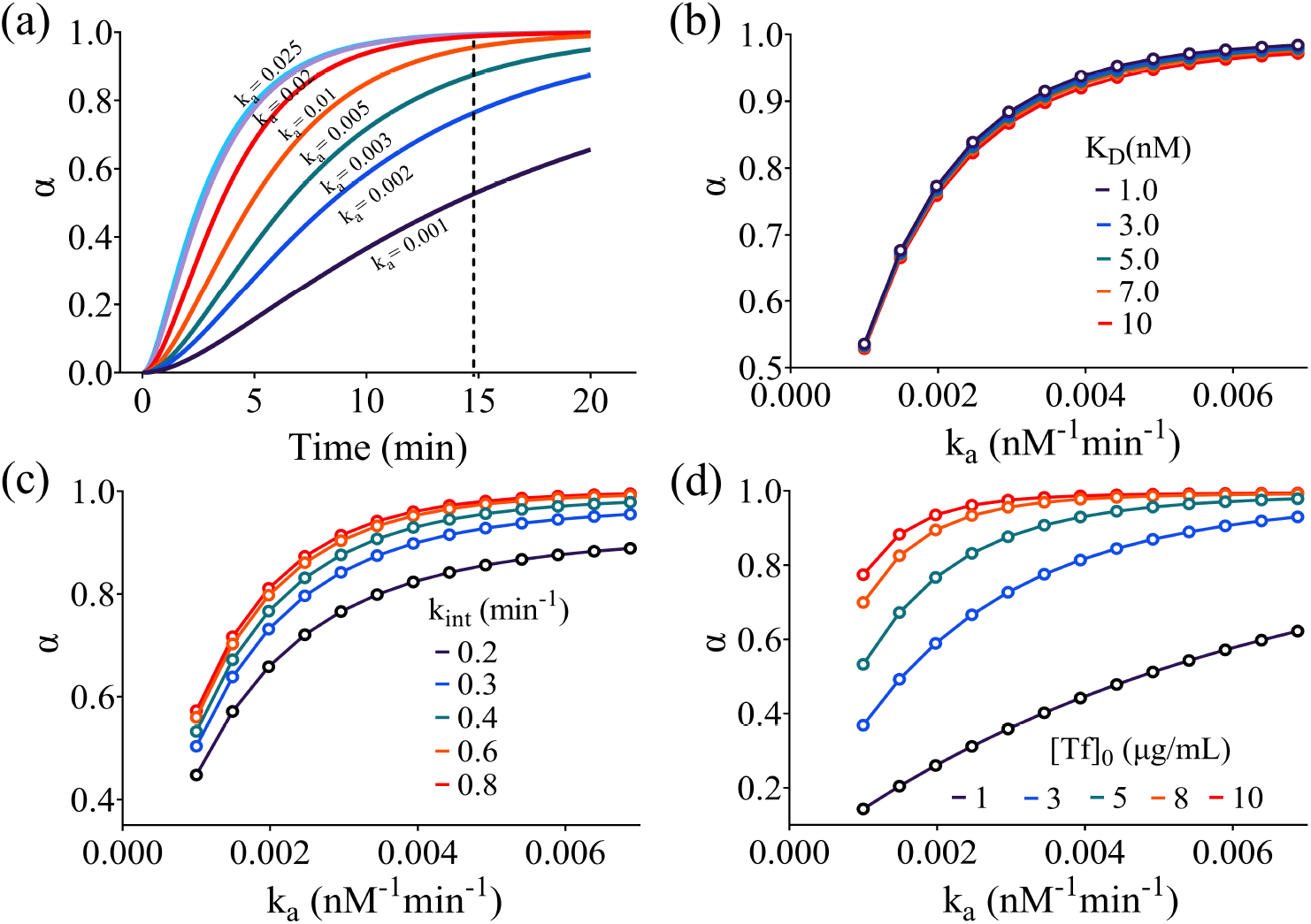
Fraction of Tf bound with TfR internalized out of total number of TfR (α). (a) as a function of time for different values of *k*_*a*_ and fixed *k*_*int*_ = 0.4 *min*^−1^, *K*_*D*_ = 5 nM, and [*T f*]_0_ = 5 µg/mL where red dashed line corresponds to 15 min; (b) at 15 min as a function of *k*_*a*_ for different values of *K*_*D*_ and fixed *k*_*int*_ = 0.4 *min*^−1^ and [*T f*]_0_ = 5 µg/mL; (c) at 15 min as a function of *k*_*a*_ for different values of *k*_*int*_ and fixed *K*_*D*_ = 5 nM and [*T f*]_0_ = 5 µg/mL; (d) at 15 min as a function of *k*_*a*_ for different values of [*T f*]_0_ and fixed *k*_*int*_ = 0.4 *min*^−1^ and *K*_*D*_ = 5 nM.

The receptor ligand complex formation is also influenced by ligand concentration in the system. So, to understand the role of Tf-A546 concentration over their internalization, in the presence and absence of active fluctuations, we choose five different concentrations of Tf-A546: 1 µg/mL, 3 µg/mL, 5 µg/mL, 8 µg/mL, and 10 µg/mL. Here, cells were incubated with different concentration of transferrin for 15 min. Our observations in **Fig. 6** show that the uptake of Tf-A546 increased with increasing concentration of Tf-A546 up to 8 µg/mL and became saturated at higher concentrations (see **Fig. 6k**). This suggests that ligand density plays an important role in Tf internalization. Comparing absence vs presence of enzymatic catalysis (5 nM urease, 100 mM urea), the Tf-A546 uptake was significantly enhanced up to a Tf-A546 concentration of 8 µg/mL with no significant change observed at higher Tf-A456 concentration **Fig. 6l**. Even at 8 µg/mL, the increase in TfA456 uptake is much smaller than those observed at lower Tf-A456 concentrations. Using the theoretical model, we can see that large fraction of TfR receptor are internalized at relatively low values of *k*_*a*_ for high Tf concentration 8 µg/mL **Fig. 5d**. Thus, at these high Tf concentrations, any enhancement of *k*_*a*_ due to higher diffusivity is presence of enzymatic catalysis will lead to marginal or no increase in Tf internalization. While the internalization of Tf is often viewed as part of constitutive endocytosis of TfR, prior work has also posited that Tf binding itself could trigger the internalization of the receptor through activation of downstream signaling pathways involved in CME (*41*). The study showed that addition of Tf to cultured epithelial cells mediates phosphorylation of two important components of the endocytic machinery, namely, the large GTPase dynamin 2 (Dyn2) and its associated actin-binding protein, cortactin (Cort) (*41*). This suggests an intriguing possibility of change in binding rate of Tf to TfR, further leading to an altered internalization rate via the intermediate signaling pathways, which can in turn influence the amount of Tf internalized. This hypothesis is in part supported by our theoretical model where we observed that change in internalization rate at a particular value of binding rate can also lead to an increase in Tf uptake, though not as large an increase as that may be observed with increasing binding rate **Fig. 5c**. Therefore, to investigate the potential role of these downstream constituents in enzymatic activity enhanced Tf internalization, we performed experiments using dynasore which is an inhibitor of dynamin (*41–43*). Cells were treated with 80 µM dynasore for 30 min. After inhibition, we incubated cells with 80 µM dynasore and different concentrations of Tf-A546 for 15 mins. An important observation to note is that even after inhibition of dynamin, which would in turn inhibit clathrin-mediated endocytosis, we still observed Tf uptake by cells **Fig. 7**. This suggests that Tf internalization is also mediated by CME-independent mechanisms. Furthermore, in contrast to previous experiments, we did not see any significant change in the fluorescence intensity per unit area (Φ) of transferrin in the presence and absence of enzyme activity (5 nM urease and 100 mM urea) **Fig. 7g**. On the other hand, consistent with prior observation, even in the presence of dynasore, we saw an increase in the Tf internalization with increasing Tf concentration. However, the observed increase in Tf uptake as a function of Tf concentration reached saturation around 8 µg/mL **Fig. 6l** in the absence of dynasore, and this saturation concentration decreased to around 3 µg/mL in the presence of dynasore **Fig. 7g**. Overall, the results indicate that, though Tf can be endocytosed by cells through either CME or CME-independent pathways (*9, 44*), the increase in Tf internalization in the presence of enzyme activity is mediated by dynamin-based CME and not due to other potential mechanisms of endocytosis.

**Figure 6:**
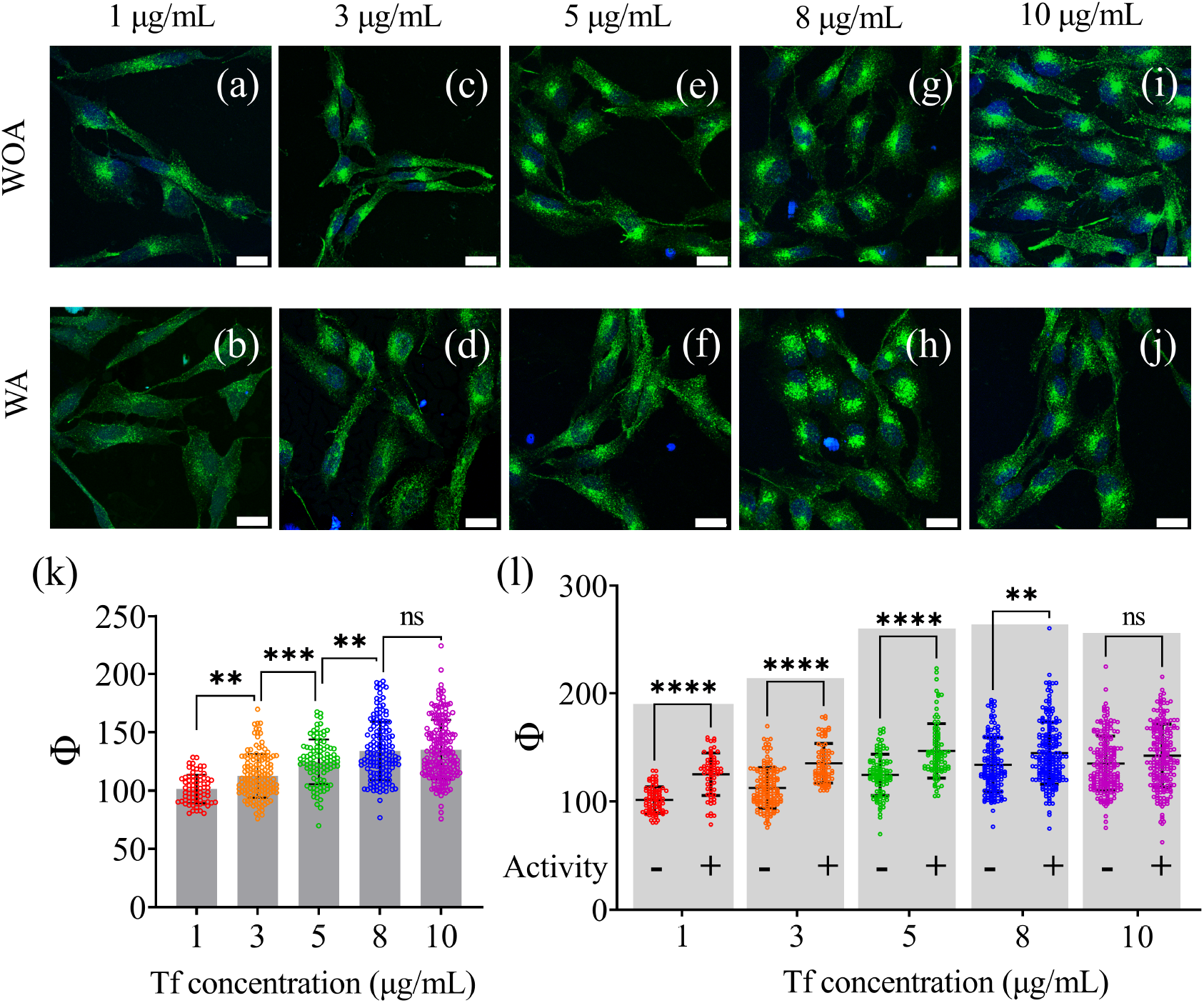
Effect of ligand concentration (Tf-A546) on endocytic internalization and the impact of enzymatic activity at each concentration. (a)–(j) Representative confocal images of RPE1 cells showing the uptake of Tf-A546 (green) without and with enzymatic (urease) catalytic activity in the extracellular media. (k) shows the variation of Φ with increasing Tf-A546 concentration in the extracellular environment. (l) Variation of Φ at different concentrations of Tf-A546 in the presence and absence of catalytic activity in the extracellular media. Scale bars represent 20 µm and error bars represent the standard errors in uptake measurements from over 80 cells. The symbols ∗∗, ∗ ∗ ∗ and ∗ ∗ ∗∗ indicate significance levels of p < 0.01, p < 0.001 and p < 0.0001 respectively, while ns denotes not significant.

**Figure 7:**
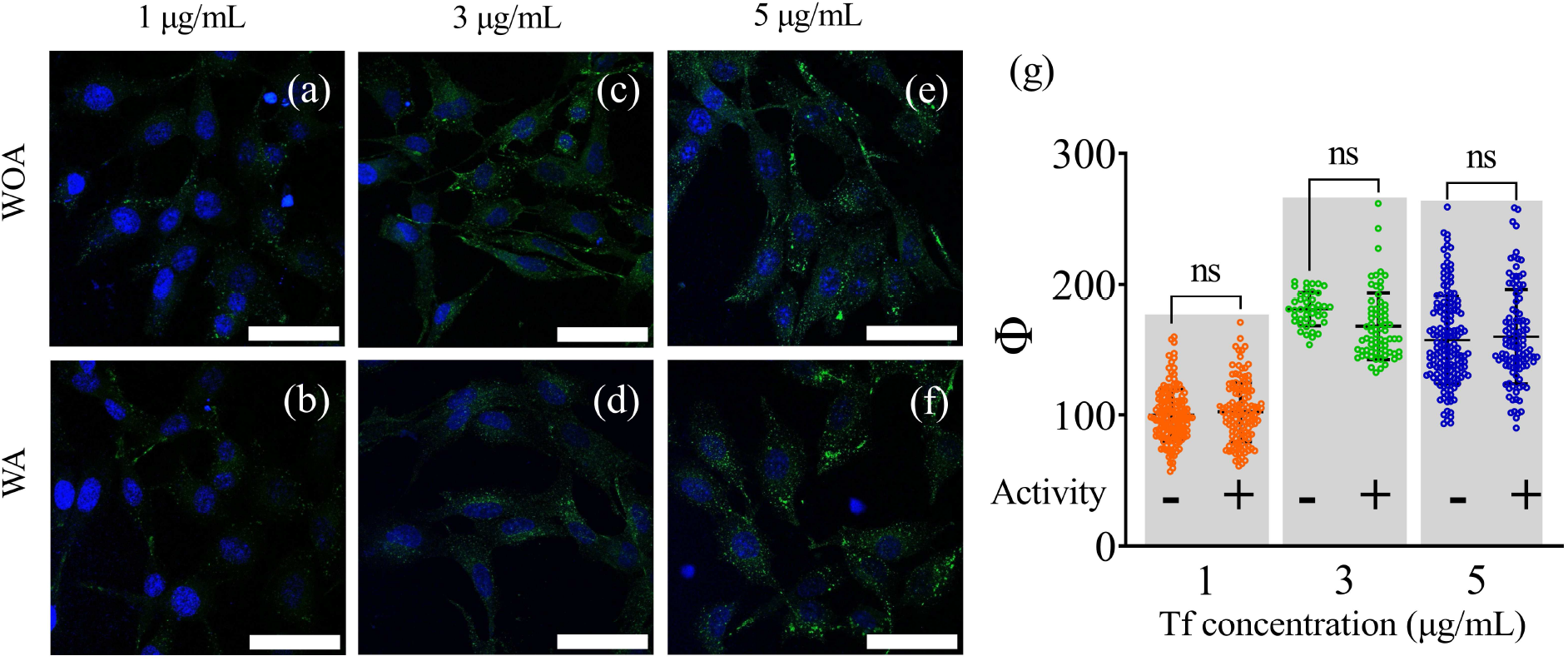
Dynamin inhibition studies for blocking clathrin mediated endocytosis of Tf-A546. (a)–(f) Representative confocal images of RPE1 cells treated with Tf and 80 µM Dynasore in the presence or absence of enzymatic (urease) catalytic activity in the extracellular media. (g) Variation of Φ at different concentrations of Tf-A546 with Dynasore, in the presence and absence of catalytic activity in the extracellular media. Scale bars represent 60 µm and error bars represent the standard errors in uptake measurements from over 40 cells. The symbol ns denotes not significant.

## Discussion

Ligand diffusion through the complex extracellular matrix and its subsequent uptake by cells are central to key biological processes including signaling, growth, and disease progression. A deeper understanding of the mechanisms that regulate ligand diffusion, receptor binding, and internalization could not only provide insights into biological organization but also facilitate the development of advanced nanosystems capable of modulating or programming biological functions. Our findings demonstrate that free enzyme activity plays a critical role in regulating molecular transport across cell membranes especially in processes such as clathrin mediated endocytosis. We observed that the diffusivity of catalytically inactive molecules (Tf-A456) increased in the presence of enzymatic reactions by as much as 40%. This increased diffusion can lead to greater collision frequency between Tf and TfR and thus, higher binding rates which can in turn mediate enhanced internalization of Tf as shown by theoretical model and experimental observations. Our results also show that while non-CME pathways can be used by cells to internalize Tf, the observed increase in Tf uptake in the presence of enzymatic activity is primarily due to CMEbased endocytosis. This can be explained by the transduction of Tf-TfR binding into downstream signaling pathways that activate cytoplasmic proteins involved in CME, particularly Src-mediated phosphorylation of dynamin. Altogether, our study suggests that increase in Tf diffusivity leads to higher binding rate for TfR, which in turn leads to increased CME-based endocytosis. This enhanced molecular and particle transport during catalytic reactions has significant implications for molecular transport processes across different barriers. This opens new avenues for developing more effective therapeutic strategies, particularly in drug delivery systems where slow diffusion is a limiting factor. The findings underscore the potential of enzyme-powered biocompatible systems in targeted drug or antidote delivery, offering promising applications in biomedical treatments.

## Acknowledgments

We thank the Central Instrument Facility at IIT Gandhinagar for confocal microscopy facility and Department of Biological Sciences and Engineering, IIT Gandhinagar for the cell culture facility. KKD thanks Anusandhan National Research Foundation (ANRF), India (CRG/2023/007588), Ministry of Education, Government of India (MoE-STARS/STARS-2/2023-0620) and IIT Gandhinagar for financial support. DB thanks SERB-DST for the Core research grant, MoE for STARS grant, Gujcost-DST and GSBTM for research funds. RR thanks the National Institutes of Health (NIGMS GM136259 and NCI CA250044) for research funds.

