## Supplementary Materials for "Enzyme-Regulated Non-Thermal Fluctuations Enhance Ligand Diffusion and Receptor-Mediated Endocytosis"

**This PDF file includes:**

Materials and Methods

Supplementary Text

Figures S1 to S5

Tables S1

### Materials and Methods

#### Cell culture

RPE1 (Retinal pigment epithelial) cell lines were grown in a T-25 flask (Nest) in Dulbecco's Modified Eagle Medium (DMEM, Sigma-Aldrich). DMEM media was supplemented with 10% fetal bovine serum (FBS; Gibco Make) and 1% penicillin/streptomycin (Pen-Strep; Gibco Make). The cells were incubated at 37 °C with 5% $CO_2$ and within 95% humidity. When the confluency reached 80%, cells were detached by using 0.05% trypsin–EDTA (Himedia) and used for further experiments.

#### Cellular uptake

For cellular uptake experiments, cells were seeded on 12 mm glass coverslips in a 24 well plate. Before experiments, cells were checked in the microscope to confirm their proper attachment and spreading. Cells were washed twice with 1× PBS and then were incubated with serum-free media, 5 $\mu$g/mL transferrin tagged with Alexa Fluor 546 (Tf-A546), 100 mM urea, and different concentrations of urease (1, 3, and 5 nM), keeping the total volume 500 $\mu$L. For the case of alkaline phosphatase enzyme, we took different concentrations of AKP (0.2, 0.4, and 0.6 nM) and 1 mM substrate p-NPP. The concentrations of the enzyme and the substrate were chosen so that the catalysis continues for the entire duration of transferrin uptake measurements. For example, with 3 nM urease and 100 mM urea, the reaction was estimated to continue for nearly a period of 27 min. The cells treated with Tf-A546, enzyme, and substrate were incubated at 37 °C for 15 min as reported in the literature. (*1*) Control experiments were performed with cells incubated with only Tf-A546. After 15 min, the cells were washed thrice with 1× PBS to remove the excess or surface-bound transferrin. The cells were then fixed by incubating in 4% paraformaldehyde for 15 min at 37 °C. This was followed by washing them three times with 1× PBS and mounting them over a glass slide using Mowiol containing Hoechst dye (Thermo Scientific), which stained the nuclei.

#### Confocal microscopy

Fixed cells were imaged using a Confocal Laser Scanning Microscope (Leica TCS SP8). The cell-attached slides were imaged using a 63× oil immersion objective. The pinhole was kept at 1

Airy unit. Different fluorophores were excited using different lasers - for Hoechst and Tf-A546, the excitation wavelengths were 405 and 556 nm, respectively. The position and shape of the nuclei were checked through Hoechst fluorescence to ensure that the cells were alive during the experiments.

#### Image analysis

Image analysis was performed manually using Fiji ImageJ software. The background from each image was subtracted first, followed by estimating the whole cell intensity using maximum intensity projection. 7-8 z-stacks were taken for each sample, and around 30 cells were used to quantify the cellular uptake.

#### Statistical analysis

Data are represented as mean ± standard deviation. Data normalization and statistical analysis were performed using GraphPad Prism 9.0. The data normalization process involved consideration of 0% as zero value and the average integrated density value of the control set as 100%, for all datasets. For statistical analysis, differences between the mean values of the control group with different test groups were calculated by using one-way ANOVA.

#### ODE model for transferrin internalization

The model was built using COPASI (version 4.44.295) and simulated in python (version 3.12.6) using the basiCO package (version 0.75). The code is available on:
GitHub:https://github.com/kparihar13/transferrin_internalization

#### Dynamin inhibition study

The endocytosis inhibition study was performed to understand the pathway by which Tf-A546 are being uptaken in presence of active fluctuations. RPE1 cells were seeded into 24-well plate on coverslips and grown until 80–90% confluency. The cells were washed with PBS and preincubated with 80 $\mu$M Dynasore in serum free DMEM media for 30 min at 37 °C. The media was decanted, and the cells were treated with the same concentration of inhibitors with Tf-A546 (1 $\mu$g/mL, 3 $\mu$g/mL, and 5 $\mu$g/mL) in serum free DMEM media. The cells were incubated at 37 °C for 15 min. The media was removed and then washed three times with glycin buffer (pH 2.2). The cells were then three time washed with PBS and fixed with 4% paraformaldehyde (PFA) at 37 °C for 15 min. They were again three times washed with PBS and mounted on the glass slide with 4′,6-diamidino-2-phenylindole (DAPI).

#### Urease activity assay

Activity assay of Jack bean urease was performed following the protocol reported in the literature. (*2, 3*) 10 $\mu$M urease solution was prepared in deionized water, and urease activity was determined by UV-vis spectroscopy. Phenol red was used as an indicator in the activity assay, which changes colour from yellow to pink with increasing pH, and corresponding absorbance was measured at 560 nm for the first 900 s. 1 nM, 3 nM, and 5 nM enzyme prepared from 10 $\mu$M stock solution, were mixed with 100 mM urea and 28.2 $\mu$M phenol red. The total volume of the experimental solution was kept fixed at 2 mL. **Fig. S1(a)** shows the measured urease kinetics curves. To calculate reaction rate, we measured the slope of the absorbance curves up to first 350 s and normalized them with the average value of minimum reaction rate data set (shown in **Fig. S1(b)**).

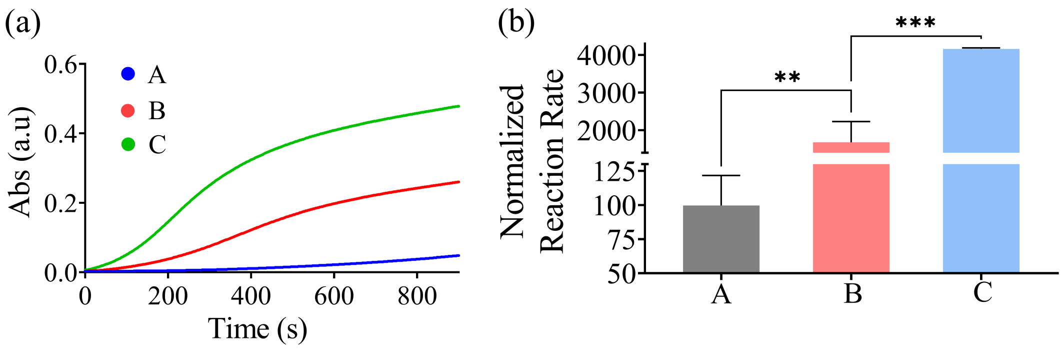


**Figure S1**: (a) Urease activity was estimated from the UV-Vis absorbance curve of Phenol red with time, measured at a wavelength of 560 nm in 100 mM aqueous urea solution. (A), (B) and (C) represent kinetic curves for 1 nM, 3 nM, and 5 nM concentrations of urease respectively. (b) Normalized reaction rates for (A) 1 nM urease, (B) 3 nM urease, and (C) 5 nM urease, each with 100 mM urea. The symbols ** and * * * indicate significance levels of $p < 0.01$ and $p < 0.001$, respectively.

### AKP activity assay

Activity assay of Alkaline Phosphatase (AKP) from bovine intestinal mucosa (Sigma) was performed following the protocol reported in the literature (*3*), with different concentrations of AKP (0.2 nM, 0.4 nM and 0.6 nM) and 1 mM p-nitro phenyl phosphate (p-NPP) (Sigma) in deionized water. The total volume of the experimental solution was kept fixed at 2 mL. AKP catalyzes the p-NPP into p-nitrophenol (p-NP) and inorganic phosphate (P) and the absorbance of p-nitrophenol was measured at 405 nm for first 600 s using UV-vis spectroscopy. By measuring the rate of product formation in the form of absorbance, we estimated the enzyme activity in solution. **Fig. S2 (a)** shows the measured AKP kinetics curves. To calculate the reaction rates, we measured the slope of the kinetic curves up to the first 150 s and normalized them with the average value of minimum reaction rate data set (shown in **Fig. S1(b)**).

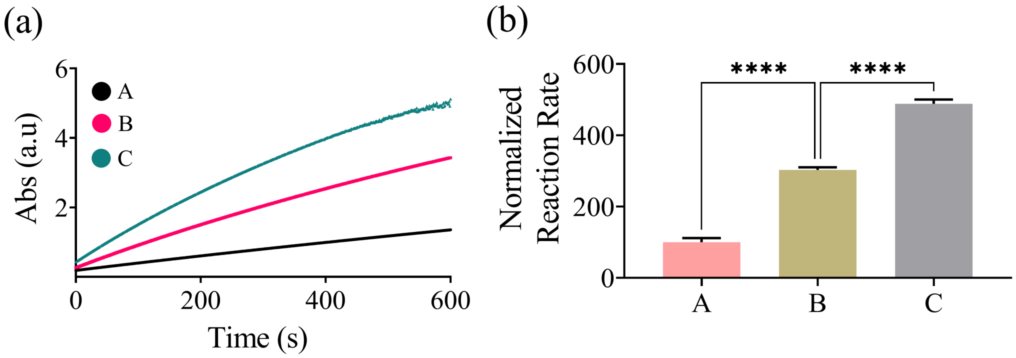


**Figure S2**: AKP activity was estimated from the rates of absorbance of p-NP with time, measured at a wavelength of 405 nm in DI water. (A), (B) and (C) represent kinetic curves for 0.2 nM AKP with 1 mM p-NPP, 0.4 nM AKP with 1 mM p-NPP, and 0.6 nM AKP with 1 mM p-NPP, respectively. (b) Normalized reaction rates for (A) 0.2 nM AKP, (B) 0.4 nM AKP, and (C) 0.6 nM AKP, each with 1 mM p-NPP. The symbols $***\,*$ indicate significance levels of $p < 0.0001$.

### Fluorescence Correlation Spectroscopy (FCS) measurements

Diffusion studies of Tf-A546 were carried out on a Confocal Fluorescence Correlation Spectrometer (Model No: HO-SP-MFCS2C). A water immersion objective 60× 1.2 NA was used to focus the excitation light of 532 nm from a diode laser onto the sample (50 $\mu$L) on a coverslip. Fluctuations in fluorescence intensity from the diffusion of transferrin molecules were auto-correlated and fit by a multicomponent 3D model to determine the diffusion coefficients of the transferrin molecules. Eq (1) defines the autocorrelation of the intensity signal.

$$G(\tau) = \frac{1}{N}\left[1 + \frac{4D\tau}{\omega_{xy}^2}\right]^{-1}\left[1 + \frac{1}{k^2}\frac{4D\tau}{\omega_{xy}^2}\right]^{-1/2} \tag{S1}$$

Here, N is the average number of fluorescent molecules in the observation volume, $\tau$ is the autocorrelation time, and k is the structure factor, which is defined as the ratio of height to width of the illumination profile (~ 3.19), and D is diffusion coefficient of the molecules crossing a circular area with radius $\omega_{xy}$ (~ 350 nm). The optical system was calibrated before each experiment using 50 nM rhodamine 6G in distilled water. Autocorrelation curves were fit to Eq.[1] using the Levenberg-Marquardt non-linear least squares regression algorithm with Origin software to determine D. The quality of the fitted curves was assessed based on chi-square ($\chi^2$) analysis.

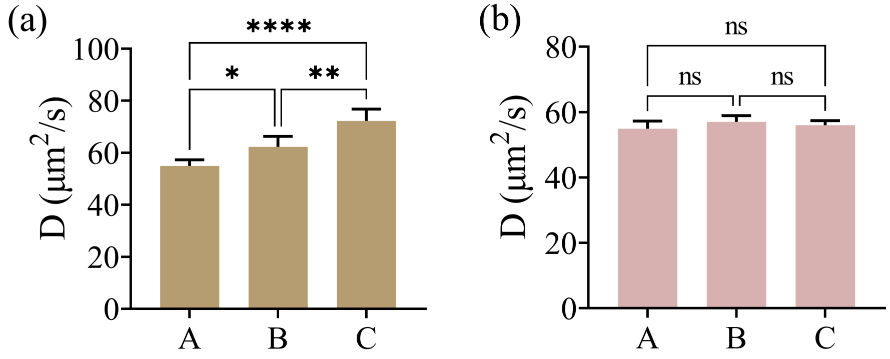


**Figure S3**: (a) Measured diffusion coefficients D of Tf-A546 in (A) DI water, (B) 3 nM urease and 100 mM urea, and (C) 5 nM urease and 100 mM urea. (b) Diffusion coefficients D of Tf-A546 in the presence of only (A) DI water, (B) 5 nM urease, and (C) 100 mM urea. Error bars represent standard deviations calculated from three independent measurements under identical conditions. The symbols $*$, $**$, and $* * **$ denote significance levels of $p < 0.05$, $p < 0.01$, and $p < 0.0001$ respectively while ns denotes not significant.

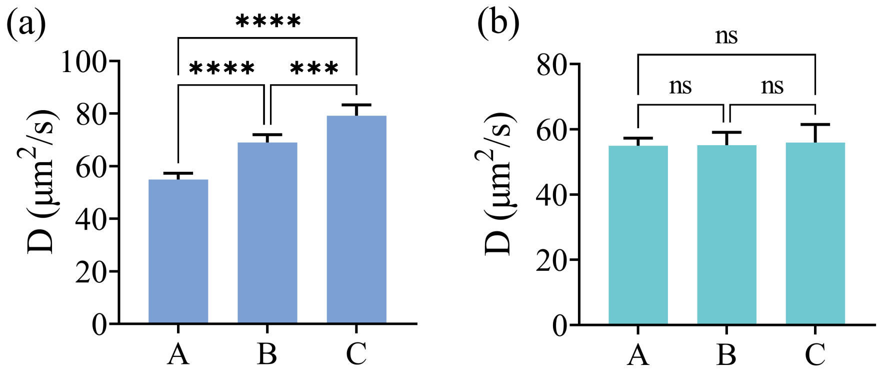


**Figure S4**: (a) Measured diffusion coefficients D of Tf-A546 in (A) DI water, (B) 0.4 nM AKP and 1 mM pNPP, and (C) 0.6 nM AKP and 1 mM pNPP. (b) Diffusion coefficients D of Tf-A546 in the presence of only (A) DI water, (B) 0.6 nM AKP, and (C) 1 mM pNPP. Error bars represent standard deviations calculated from three independent measurements under identical conditions. The symbols $***$ and $****$ denote significance levels of $p < 0.001$ and $p < 0.0001$, respectively, while ns denotes not significant.

### Uptake of fluorescence labelled urease

In our experiment, we considered that most of the urease molecules are present in the extracellular environment and not uptaken by the cells during the incubation time. To conform this, we tagged urease with a fluorescent dye and incubated the cells with the labelled urease. After incubation, we imaged the cells to quantify the uptake of urease. First for tagging the enzyme, Jack bean urease (type C-3; Sigma-Aldrich) was tagged with a thiol-reactive dye, Dylight 549 (ex/em: 549/568; Thermo Fisher Scientific). The reaction of the fluorescent probe (40 $\mu$M) with urease (2 $\mu$M) was carried out in 150 mM phosphate buffer (pH 7) at room temperature for 4-6 h under gentle stirring as mentioned in earlier studies. (*4*) The enzyme-dye complexes were further purified using membrane dialysis (100 kDa pores; Amicon ultra-4 centrifugal filter unit, Millipore) to reduce free-dye concentration. After that cells were incubated with dye tagged urease for 15 min, which was our experiment time. Urease uptake was quantified by confocal imaging, where we did not observe any significant change in the fluorescence intensity of the cells treated with tagged urease compared to those without urease treatment.

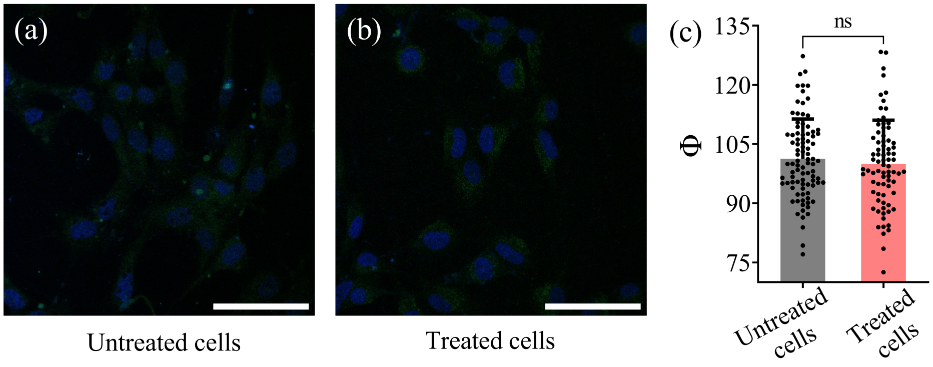


**Figure S5**: Confocal images of RPE cells (a) without and (b) with dye tagged urease treatment. (c) Normalized fluorescence intensity per unit area (Φ) of cells, measured for cells treated and untreated with fluorescently tagged urease. Scale bar represents 60 µm, and error bars represent the standard errors in uptake measurements from over 100 cells. The symbol ns denotes not significant.

### ODE model for transferrin internalization

We model the uptake of Tf using an ordinary differential equation (ODE) model. The model consists of 2 reactions and 4 species, namely Tf, Tf receptors (TfR), receptor bound Tf (Tf-TfR), and internalized Tf-TfR (Tf-TfR$_{int}$). We assume that TfR is internalized only when bound to Tf. **Table 1** describes the reactions and range of values that have been reported for the binding rate of Tf with TfR ($k_a$), dissociation constant for Tf-TfR ($K_D$), and internalization rate for Tf-TfR ($k_{int}$).

**Table S1**: **Reactions in the ODE model and range of values reported in literature for their corresponding rate constants**

| Reactions | Rate Constants | References |
| --- | --- | --- |
| $Tf + TfR \Leftrightarrow Tf - TfR$ | $k_a = [0.001 - 0.025] nM^{-1} min^{-1}$ | [ (*5*), (*6*), (*7*), (*8*)] |
| | $k_d$ = dissociation rate = $K_D \cdot k_a min^{-1}$ | |
| | $K_D$= [1-10] nM | [ (*9*), (*10*)] |
| $Tf + TfR \Leftrightarrow Tf - TfR_{int}$ | $k_{int} = [0.2 - 0.8] min^{-1}$ | [ (*5*), (*6*), (*7*), (*8*)] |

Prior studies have estimated the cycling time for Tf to be 16.9 min (*5*), 14 min (*7*), and 22.7 min (*11*). Based on these prior experimental estimates, we incubated cells with Tf-A456 in our experiments for 15 min, to minimize recycling of internalized Tf-A456. Thus, in the construction of our ODE model, we assume that the recycling of Tf to be negligible and do not include a recycling reaction. The initial concentrations for Tf-TfR and Tf-TfR$_{int}$ were assumed to be 0. The initial concentration for Tf was based on the amount of Tf ($\mu$g/mL) used in the experiments. For 5 $\mu$g/mL of Tf-A456, the initial concentration would thus be

$$[Tf]_0 = \left( \frac{(5\,\mu g/\text{mL})\left(10^{-6}\,g/\mu g\right)\left(10^{3}\,\text{mL/L}\right)}{80000\,g/\text{mol}} \right) \times 10^{9}\,\text{nmol/mol} = 62.5\,\text{nM}$$

Number of Tf receptors per cell have been previously reported to be of the order of $10^5$ [9, 10]. In our experiments, we calculated approximately 1,500 cells on the coverslip, and the solution volume used was 500 $\mu$L, the initial concentration for TfR was calculated as

$$[TfR]_0 = \left( \frac{1500 \times 10^{5}}{\left(6.023 \times 10^{23}\,\text{mol}^{-1}\right)\left(500 \times 10^{-6}\,L\right)} \right) \times 10^{9}\,\text{nmol/mol} = 5 \times 10^{-4}\,nM$$
